# Reduced cerebellar Gq-protein signaling elicits early onset spatial navigation deficits in a SCA6 mouse model

**DOI:** 10.1101/2023.09.12.557443

**Authors:** Michelle Grömmke, Hannah Schulte, Candy D. C. Theis, Lena Nonnweiler, Melanie D. Mark

## Abstract

Spinocerebellar ataxia type 6 (SCA6) is a hereditary neurodegenerative disease that manifests in a late onset and progressive impairment of motor coordination, balance and speech as well as cerebellar and brainstem atrophy. It is caused by a polyglutamine expansion in the *CACNA1A* gene which bicistronically encodes the α1A-subunit of the P/Q-type voltage-gated calcium channel and the transcription factor α1ACT. To date, no effective treatment exists and the exact pathobiology is controversially discussed; especially the impact on cognition is poorly understood. Here, we demonstrate that SCA6 84Q mice exhibit cognitive deficits in their spatial navigation abilities. Surprisingly, spatial memory impairments develop prior to motor impairments at 5 months of age. By expressing and stimulating a Gq-protein coupled designer receptor exclusively activated by a designer drug (Gq-DREADD) in the cerebellum, we were able to counteract these spatial navigation deficits indicating that a reduced Gq-protein signaling is part of the SCA6 phenotype. Electrophysiological recordings in anaesthetized mice further revealed that Purkinje cells (PCs) of SCA6 84Q mice exhibit a disrupted spontaneous simple spike activity that precedes the development of both cognitive and motor deficits. Concurrently, PC dysfunction was further confirmed by elevated numbers of torpedoes found in the proximal axon of PCs throughout the cerebellum. Overall, our study raises awareness to survey cognitive abnormalities more carefully during clinical examination to detect the disease earlier and potentially optimize the individual treatment by enhancing PC signaling.

**Significance statement:** SCA6 is a hereditary neurological disease that is mainly characterized by the late-onset development of progressive motor deficits. Here we show, using a SCA6 mouse model, that cognitive impairments in spatial navigation are also a non-negligible feature of the disease which manifests earlier than the motor deficits. Moreover, we demonstrate that these spatial navigation deficits are caused by a reduced Gq-protein signaling in the cerebellum. Electrophysiological and histological analysis further confirmed dysfunctional PC signaling even before the onset of first symptoms. Since no effective treatment is available for SCA6 patients, early onset stimulation of PC signaling may be a new therapeutic approach.

## Introduction

SCA6 is a neurodegenerative disease caused by a CAG repeat expansion in exon 47 of the *CACNA1A* gene (Zhuchenko et al., 1997) which bicistronically encodes the pore-forming α1A-subunit of the P/Q-type Ca^2+^-channel and the transcription factor α1ACT (Ophoff et al., 1996; Du et al., 2013). The latter modulates the expression of genes associated with cerebellar and PC development and mediates neurite outgrowth (Du et al., 2013). The P/Q-type channel, on the other hand, is a high-voltage-gated calcium channel which is primarily involved in synaptic transmission but also plays a pivotal role in modulating membrane excitability and synaptic plasticity (Mintz et al., 1995; Catterall and Few, 2008). The *CACNA1A* gene is expressed throughout the whole brain, but because of its strong expression in cerebellar PC and granule cells (GrCs) (Westenbroek et al., 1995; Kulik et al., 2004), SCA6 symptoms are thought to relate exclusively to cerebellar dysfunctions including gait ataxia, upper limb incoordination and intention tremor as well as dysarthria, nystgamus and diplopia (Watase et al., 2008; Mark et al., 2015; Du and Gomez, 2018). Additionally, loss of PCs and GrCs (Sasaki et al., 1998) and morphological anomalies of PCs like heterotopic nuclei, somatic sprouts and axonal swellings have been described (Yang et al., 2000). Since there is currently no treatment and the pathobiology is controversial, it is crucial to gain further knowledge of the mechanisms underlying SCA6. In particular, the impact of SCA6 on cognitive processes is not completely understood. While no severe cognitive dysfunctions were found in a study conducted by Globas et al. (2003), mild decreases in verbal fluency as well as impairments in visual-motor processing, executive functions and social cognition were described (Suenaga et al., 2008; Pereira et al., 2017; Giocondo and Curcio, 2018; Abdelgabar et al., 2019). Mouse models are widely used to study the unique cellular features underlying SCA6 development and pathology. CT-long^Q27^ mice which express elongated C-terminal fragments of the P/Q-type channel develop typical SCA6 symptoms including ataxia and PC degeneration, highlighting the importance of the C-terminus in SCA6 pathology (Mark et al., 2015). Recently, these mice were shown to exhibit a decrease in innate fear and defense behavior (Bohne et al., 2022). Another rodent model that closely mimics SCA6 both genetically and phenotypically are SCA6 84Q mice (Watase et al., 2008). SCA6 84Q mice express the P/Q-type Ca^2+^-channel carrying a hyperexpanded (84Q) C-terminus and show a midlife onset of motor impairments at 7 months accompanied by PC firing deficits while showing no change in P/Q-type channel conductance or kinetics (Watase et al., 2008; Jayabal et al., 2016). Morphological abnormalities develop at a late stage of the disease and manifest as neuronal inclusions in the PC cytoplasm at 22 months as well as PC cell loss and enhanced axonal swellings after 2 years (Watase et al., 2008; Jayabal et al., 2015; Ljungberg et al., 2016).

To determine if SCA6 interferes with cognition, we tested SCA6 84Q mice in two spatial navigation paradigms and found strong spatial memory deficits developing at 5 months of age in the Morris water maze (MWM) and only mild impairments in the starmaze. Similar results were described for L7-PKCI (pseudosubstrate PKC inhibitor under the control of the PC specific promoter L7) mice which exhibited diminished long-term depression (LTD) at the parallel fiber-PC (PF-PC) synapse (Zeeuw et al., 1998; Burguière et al., 2005). Likewise, impaired LTD at the PF-PC synapse was found in another SCA6 mouse model (Mark et al., 2015). Therefore we aimed to promote LTD by enhancing Gq-protein signaling, which is tremendously involved in LTD-induction (Hartmann et al., 2004), using a Gq-DREADD (Armbruster et al., 2007) and were able to improve the spatial navigation performance of SCA6 84Q mice. To further examine if PC dysfunction may contribute to the observed phenotype, we conducted a detailed electrophysiological and histological analysis and found clear firing deficits and elevated numbers of torpedoes throughout all ages tested starting at 3 months.

## Materials and methods

### Animals

For all experiments male and female homozygous SCA6^84Q/84Q^/Cas9^+/+^/Pcp2-cre mice, from now on shortly referred to as SCA6 84Q obtained by crossing SCA6^84Q/84Q^ mice (JAX stock #008683, (Watase et al., 2008), Cas9^+/+^ mice (JAX stock #024857, (Platt et al., 2014) and Pcp2-cre mice (JAX stock #004146, (Barski et al., 2000)) were used. Cas9^+/+^/Pcp2-cre mice of both sexes were used as control animals throughout experiments and are therefore referred to as controls. Mice were housed under a twelve hour day and night cycle in individually ventilated cages with unlimited access to food and water. Standard PCR analysis was used to validate transgene expression: SCA6 84Q forward 5’-ACGTGTCCTATTCCCCTGTGATCC-3’, SCA6 84Q reverse 5’-ACCAGTCGTCCTCGCTCTC-3’, Cre forward 5’-TCTCACGTACTGACGGTGG-3’, Cre reverse 5’-ACCAGCTTGCATGATCTCC-3’. Genotyping for Cas9 transgene was performed with primer sequences provided by Jackson laboratories. All experiments were performed with approval of a local ethics committee (Bezirksamt Arnsberg) and the animal care committee of North Rhine-Westphalia (LANUV; Landesamt für Umweltschutz, Naturschutz und Verbraucherschutz Nordrhein-Westfalen, Germany). The study was carried out in accordance with the European Communities Council Directive of 2010 (2010/63/EU) for care of laboratory animals and supervised by the animal welfare commission of the Ruhr-University Bochum.

### Cloning of recombinant pAAV vectors

For all constructs the pAAV-CW3SL-EGFP vector (made available by Bong-Kiun Kaang, Addgene plasmid #61463) was used which allows higher packaging capacities and enhanced transgene expression (Choi et al., 2014). hM3D(Gq)-mCherry (made available by Bryan Roth, Addgene plasmid #50476) or control mCherry were subcloned using the InFusion HD cloning kit (Takara Bio USA). Furthermore, the CaMKIIa promoter was replaced with the CMV promoter to obtain the final pAAV-CMV-hM3D(Gq)-mCherry-CW3SL and pAAV-CMV-mCherry-CW3SL plasmids.

### Virus production

Recombinant adeno-associated viruses (AAVs) with serotype 8 were produced using the AAV helper-free system (Agilent Technologies). Human embryonic kidney (HEK) tsA201 cells were co-transfected with the recombinant pAAV vector encoding the gene of interest (30 µg), the pHelper plasmid (60 µg) carrying the adenovirus genes *VA*, *E2A* and *E4* and the pAAV RC8 plasmid (50 µg) containing the adenoviral *rep* and *cap8* genes using polyethylenimine (7.74 µM in Dulbecco’s modified Eagle’s medium, Sigma-Aldrich D6429). 72 hours post transfection the cells and the medium were collected and centrifuged at slow speed (5 min, 1500 rpm, 4°C). The supernatant was incubated with 10% (w/v) polyethylenglykol (40% PEG-800, 2.5 M NaCl) for 2 hours on a horizontal shaker at 4°C. The cell pellet was resuspended in 10 ml lysis buffer (150 mM NaCl, 50 mM Tris-HCl pH 8,5), lysed via 5-7 freeze-thaw cycles and incubated with 0.13 mg/ml DNase I and 13 mM MgCl_2_ at 37°C for 30 min. Both the supernatant and the cell suspension were centrifuged at 3700 x g for 20 min at 4°C and the PEG-precipitated pellet was resuspended in the supernatant from the cell suspension. The virus was precipitated with 10% (w/v) PEG as described earlier and resuspended in 50 mM HEPES buffer solution. For purification the virus was thoroughly mixed with an equal volume of chloroform and centrifuged at slow speed (5 min, 370 x g, room temperature). The supernatant was carefully collected, filtered with a 0.22 µm membrane and precipitated with 10% (w/v) PEG. The virus was resuspended in a minimal volume of phosphate-buffered saline (PBS) containing 0.001% Pluronic F-68 and stored at −80°C.

### Virus injection

For injection of the recombinant AAVs mice were anesthetized with isoflurane (initial anesthesia at 5% in 1.1 L/min air and maintenance at 1.5-2% in 1.1 L/min air), placed on a 37°C heating plate and fixed in a stereotactic frame (SR-6M-HT, Narishige). Carprofen (2 mg/kg) was administered subcutaneously for analgesia. The skull was exposed via incision of the skin along the midline and lidocaine was applied for additional local anesthesia. Three small craniotomies above the cerebellum (AP: −6.5, ML: −1, 0 and +1) were drilled into the skull allowing access to the brain tissue. A custom-made glass pipette filled with 1 µl of the respective virus was lowered into the brain with a micromanipulator (SM-12, Narishige) to −1800 µm and the virus was injected in small amounts in 200 µm steps until −400 µm via pressure injection. Mice received analgesia via subcutaneous injection of carprofen (2 mg/kg) on the first 4 days, were monitored daily and housed individually and had a 14 day recovery for post-surgical care.

### Spatial navigation tests

To assess cognitive abilities mice were tested in the MWM and star maze. In both tests, mice had to find a submerged hidden platform (diameter 10 cm) in a tank (diameter 96 cm) filled with 26 ± 1°C opaque water. For opacity the water was thoroughly mixed with coffee creamer (∼2 kg depending on the brand). While the MWM is an aquatic open field, the starmaze consists of five potential target alleys radiating from a central pentagonal ring creating restricted swimming trajectories (Morris, 1984; Burguière et al., 2005). For orientation four different visual cues at each compass direction were placed around the water tank. A white curtain surrounded the arena to reduce external distraction. To minimize stress mice were handled 5 min per day for 5 days prior to the test to become accustomed to the experimenter. For 10 (MWM) or 7 (starmaze) consecutive days mice performed 4 trials/day with an intertrial interval of 120 sec. In the MWM mice were released into the water at one of four starting positions in a pseudorandomized order with comparable swimming distances to the hidden platform (Vorhees et al., 2004). The platform was located at a fixed position 10 cm from the edge. In the starmaze the platform was located in one, fixed target alley while the other four target alleys were pseudorandomly used as starting points. Individual trials were completed when the mice reached and stayed on the platform for 2 sec or when 90 sec passed. On the first training day if a mouse did not find the platform within 90 sec in all 4 trials, it was gently placed on the platform until it learned to stay on the platform. During Gq-DREADD experiments, the DREADD agonist clozapine N-oxide (CNO, Hello Bio HB6149) was administered intraperitoneally at a concentration of 1 mg/kg 30 min prior to the experiment. For data acquisition the EthoVision XT 11.5 (Noldus Information Technology) software was used and the escape latency, total distance moved, swimming velocity and mean distance to the platform were analyzed. To determine the efficacy of spatial navigation in the MWM each trial was assigned to 1 of 9 different search patterns based on (Brody and Holtzman, 2006) for predefined training days 1, 4, 7 and 10. Additionally, for the starmaze a custom made script implemented in Matlab (MathWorks) was used to generate three-dimensional heatmaps of the swimming paths.

### Electrophysiological *in vivo* recordings

Extracellular electrophysiological recordings of PCs were made as previously described (Mark et al., 2015). Briefly, mice were anesthetized with isoflurane (initial anesthesia at 5% in 1.1 L/min air and maintenance at 1.5-2% in 1.1 L/min air) and placed on a 37° C heating plate. The head was fixed using a stereotactic frame (SR-6M-HT, Narishige) and an incision was made along the midline to expose the skull. A craniotomy was made above lobules IV/V and VI of the cerebellar vermis and the *dura mater* was removed. A seven-channel multi electrode system (Eckhorn, Thomas Recording) was placed perpendicularly above the craniotomy with loose contact to the brain surface and platinum-tungsten electrodes with a resistance of 2-3 mOhm at 1 kHz (Thomas Recording) were lowered individually into the brain tissue. Signals from each channel were amplified and band-pass filtered at 0.1-8 kHz with a multichannel signal conditioner (CyberAmp380, Axon Instruments) and further sampled with 32 Hz using an A/D converter (NI PCI-6259, National Instruments). Custom made programs implemented in Matlab (MathWorks) were used to monitor the signals in real time and to analyze the data. The recorded simple spikes were sorted offline on the basis of the action potential amplitude and form. The mean spike rate and predominant spike rate (= 1 / median interspike interval) as well as the coefficient of variation for interspike intervals (ISI) CV (= standard deviation (ISI) / mean (ISI)) and the coefficient of variation for adjacent intervals of ISIs CV2 (= 2 | ISI_n+1_ – ISI_n_ | / (ISI_n_ + ISI_n+1_)) (Holt et al., 1996; Shin et al., 2007) as indicators for firing precision were calculated for each cell.

### Histology

For histological analysis mice were deeply anesthetized with ketamine (100 mg/kg) and xylazine (10 mg/kg) and transcardially perfused with PBS and 4% paraformaldehyde (in PBS, pH 7.2-7.4). Brains were removed, post-fixed for maximal 24 hours and cryoprotected in 30% (w/v) sucrose (in PBS). Frozen brain slices were made with a cryomicrotome (Leica CM3050 S). To validate proper virus expression 35 µm thick coronal sections of the cerebellum were collected on microscopic slides and directly embedded in Mowiol 4-88 supplemented with DABCO (25 mg/ml). To determine the density of axonal swellings in PCs 45 µm thick sagittal slices were collected in Tris-buffered saline (TBS) and immunohistochemically stained. Therefore, slices were heated in sodium-citrate buffer (pH 6) at 80°C for 30 min for antigen retrieval and blocked in TBS containing 0.5% Triton X-100 and 10% normal donkey serum (NDS) for two hours at room temperature. Slices were incubated overnight with the primary antibodies mouse anti calbindin D28K (1:1000, Sigma-Aldrich C9848) and rabbit anti IP3 receptor (1:1000, Abcam ab5804) in TBS containing 0.5% Triton X-100 and 1% NDS. Secondary antibodies donkey anti mouse conjugated with CF^TM^ 555 (1:1000, Sigma-Aldrich SAB4600060) and donkey anti rabbit conjugated with DyLight 650 (1:1000, Invitrogen SA5-10041) in TBS containing 0.5% Triton X-100 and 1% NDS were incubated with the slices for two hours at room temperature. Slices were transferred to microscopic slides and again embedded in Mowiol 4-88 supplemented with DABCO (25 mg/ml). An inverted confocal laser scanning microscope (Leica, TCS SP5 II) was used to image the brain slices. For torpedo count vermal lobules 3 and 8 as well as Crus I and Crus II of the hemispheres were imaged. The number of calbindin- and IP3-receptor-positive axonal swellings with a width ≥3 µm was determined and normalized as the number of torpedoes per 100 µm using ImageJ (v1.53t, NIH) software (Ljungberg et al., 2016; Louis et al., 2019).

### Statistics

When applicable, data are presented as mean ± standard error of the mean (SEM). Statistical significance was calculated with a significance level p<0.05 using the SPSS Statistics software (IBM). For the spatial navigation paradigms learning was compared using a two-way ANOVA with repeated measures followed by pairwise comparisons. For electrophysiological and histological data, a two-way ANOVA with subsequent pairwise comparisons were employed. Detailed information about the statistics for each figure are summarized in the extended data, tables S1-S5 and are referred to in the figure legends.

## Results

### SCA6 84Q mice develop early-onset spatial memory deficits

For several SCAs cognitive decline has been described to be part of the disease pathology (Ma et al., 2014). However, SCA6 is supposed to exclusively affect motor behavior and the connection to cognitive impairments remains critically reviewed (Globas et al., 2003; Suenaga et al., 2008; Pereira et al., 2017; Bohne et al., 2022). SCA6 84Q mice were shown to develop midlife motor deficits at 7 months of age (Watase et al., 2008; Jayabal et al., 2015), whereas the impact of the polyQ mutation on cognitive abilities of these mice is unknown. To asses if SCA6 84Q mice develop spatial memory deficits as a sign for cognitive decline during disease progression, we tested these mice in two aquatic spatial navigation paradigms, the MWM (Fig. 1A) and starmaze (Fig. 2A) at 3, 5, 7 and 12 months of age.

**Figure 1:**
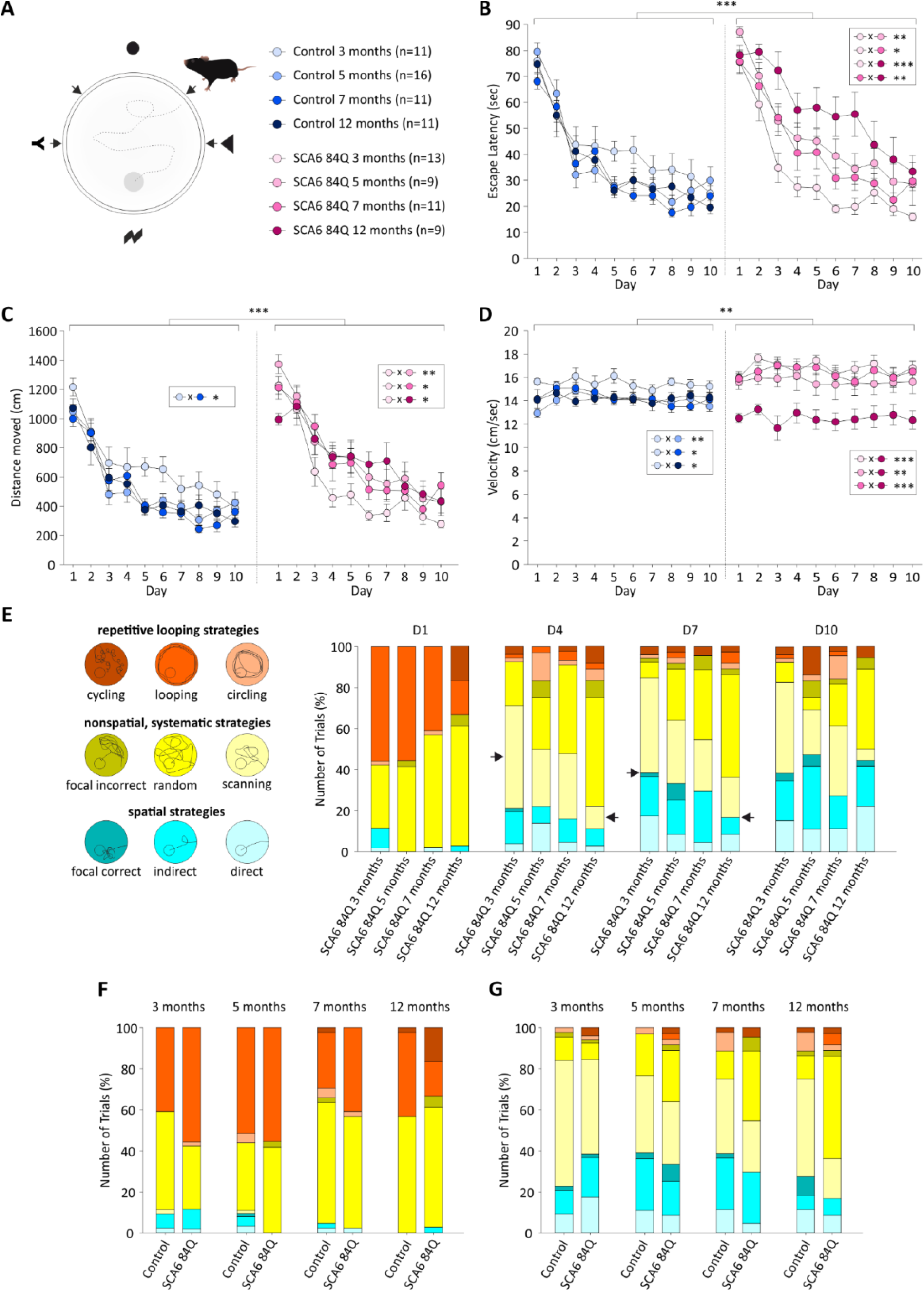
SCA6 84Q mice develop spatial memory impairments in the MWM. **(A)** Scheme of the MWM. Mice searched for a hidden platform in an aquatic open field using external visual cues (circle, triangle, lightning, Y) located at the 4 cardinal directions. Mice were trained for 10 consecutive days consisting of 4 trials/day from 4 pseudorandomized starting positions (marked by arrows). Both SCA6 84Q and control mice learned the task during the 10 days as indicated by a continuous decrease in the **(B)** escape latency and **(C)** overall distance moved (p<0.001 for both). However, compared to controls, SCA6 84Q mice exhibited higher escape latencies (p<0.001) and longer distances moved (p<0.001), indicating that SCA6 84Q mice manifested spatial navigation deficits in the MWM at 5 months of age. **(D)** The swimming velocity measured as a parameter for potential motor deficits also differs between the two groups (p=0.009). Until 7 months of age SCA6 84Q mice swam faster than controls. A substantial reduction in swimming velocity was observed in 12 months old SCA6 84Q mice probably due to acute swimming disabilities caused by the disease progression. **(E)** Evaluation of the navigation strategies from SCA6 84Q swimming paths on days 1, 4, 7 and 10. Each trial was assigned to 1 of 9 different search strategies classified into 3 superior categories: spatial strategies, systematic but nonspatial strategies and repetitive looping strategies. In general, over the 10 day test period a shift from repetitive looping strategies towards spatial strategies was observed indicating successful learning. Age-dependent differences (highlights marked with arrows) were most prominent within the nonspatial, systematic category showing a switch from random searching to scanning behavior in younger mice while older mice tend towards more ineffective random searching. SCA6 84Q mice and their age-matched controls displayed comparable search strategies on day 1 **(F),** while differences between the two groups can best be seen on day 7 **(G)**. When applicable data are presented as mean ± SEM. Statistical significance was determined with a two-way ANOVA with repeated measures (*p≤0.05, **p≤0.001, ***p≤0.001). See also extended Table S1.

**Figure 2:**
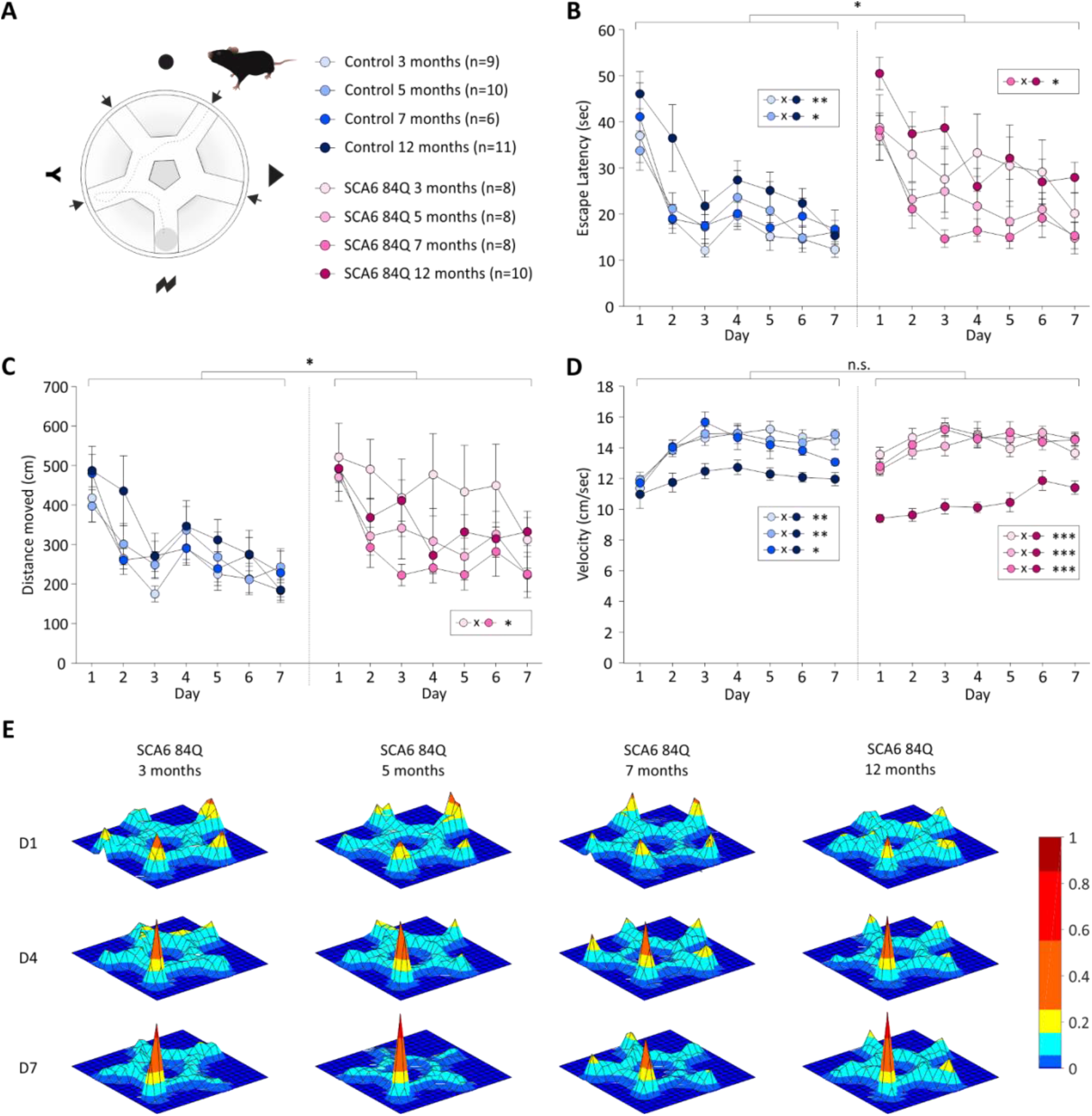
SCA6 84Q mice exhibit mild spatial navigation deficits in the star maze. **(A)** Schematic overview of the star maze. Mice searched for a hidden, submerged platform in an aquatic maze defined by restricted swimming trajectories (five potential target alleys radiating from a central pentagonal ring) with external visual cues (circle, triangle, lightning, Y) located at the 4 cardinal directions. SCA6 84Q and control mice were trained for 7 consecutive days, consisting of 4 trials/day with 4 pseudorandomized start positions (marked by arrows). Both groups successfully learned to find the hidden platform over the 7 day test period as demonstrated by a constant decline in the **(B)** escape latency and **(C)** overall distance moved (p<0.001 for both). SCA6 84Q mice displayed mild spatial navigation impairments as indicated by increased escape latencies (p=0.035) and longer distances travelled (p=0.012) compared to control mice which were not age-dependent or progressive. **(D)** SCA6 84Q and control mice navigated through the maze with comparable velocities (p=0.559). However, within both genotypes 12 months old mice swam significantly slower compared to the other age groups. **(E)** Cumulative 3D heat maps of selected days 1, 4 and 7 further illustrate comparable swimming patterns of SCA6 84Q mice at all ages tested. Over the test period mice developed a goal-directed behavior. When applicable data are presented as mean ± SEM. Statistical significance was determined with a two-way ANOVA with repeated measures (*p≤0.05, **p≤0.001, ***p≤0.001). See also extended Table S2.

First, we found that SCA6 84Q mice as well as respective control mice successfully learned to find the platform in the MWM, demonstrated by a continuous decrease in the escape latency (p<0.001 for both genotypes, ANOVA, Fig. 1B) and the total distance moved (p<0.001 for both genotypes, ANOVA, Fig. 1C) over the 10 day test period. However, SCA6 84Q mice showed a reduced performance in the MWM compared to control mice as shown by higher latencies to find the platform and total distances moved (p<0.001 for both parameters, ANOVA). Spatial memory deficits manifested at 5 months of age in SCA6 84Q mice as exhibited by their higher escape latencies (p≤0.049, pairwise comparisons) and longer distances traveled (p≤0.027, pairwise comparisons). SCA6 84Q mice also demonstrated differences in their swimming velocities (p=0.009, ANOVA) compared to control mice. Interestingly until 7 months of age SCA6 84Q mice swam faster or equally as fast as their controls indicating normal swimming abilities. A marked reduction in swimming speed was found in 12 months old SCA6 84Q mice (p≤0.009, pairwise comparisons) possibly due to acute swimming disabilities from their progressive ataxia. To evaluate the efficacy of searching strategies, individual swimming paths of SCA6 84Q mice were assigned to one of the following categories: spatial (strategic), nonspatial but systematic (strategic with low efficiency) and repetitive looping (nonstrategic) (Fig. 1E). Successful learning of the paradigm is illustrated by an increase of spatial strategies and a decrease of repetitive looping strategies. On the first day SCA6 84Q of all ages employed comparable searching strategies. Similar strategies were also found in comparison to their respective controls on day 1 (Fig. 1F). Differences between the different ages were most prominent on day 4 and 7, where we observed an age dependent decline in spatial strategies and rise in nonstrategic strategies such as nonspatial and repetitive looping. Furthermore, starting on day 4 an age-dependent shift from scanning behavior towards random swimming could be observed within the nonspatial, systematic category. Since the differences were most striking on day 7, we also compared SCA6 84Q mice with their age-matched controls on day 7 (Fig. 1G). Even though SCA6 84Q mice performed better at 3 months of age depicted by an increased number of spatial strategies, starting at 5 months of age they showed a progressive reduction in performance compared to controls, illustrated by decreasing numbers of spatial strategies and an increased random searching behavior.

Similar to the MWM, both SCA6 84Q and control groups successfully learned the starmaze task as shown by a gradual decrease in the escape latency (p<0.001 for both genotypes, ANOVA, Fig. 2B) and the total distance travelled (p<0.001 for both genotypes, ANOVA, Fig. 2C). However, unlike the results found for the MWM, we only found mild spatial memory deficits for SCA6 84Q mice in the starmaze (escape latency: p=0.035, distance moved: p=0.012, ANOVA). Additionally, we observed that both SCA6 84Q and control mice took longer to find the hidden platform at 12 months of age (control: p≤0.029, SCA6 84Q: p=0.039, pairwise comparisons), whereas no such age-dependency was observed for the total distance moved. Interestingly, learning behavior was also reflected in an increase in the swimming speed over time (p<0.001 for both genotypes, ANOVA, Fig. 1D). However, this is most likely associated to motor learning improvements over the seven training days. In this context no differences between SCA6 84Q and control mice (p=0.559, ANOVA) were found, but at 12 months of age both groups swam at a slower speed compared to respective younger mice (control: p≤0.039, SCA6 84Q: p<0.001, pairwise comparisons) which explains the increased escape latencies with no longer distances moved at 12 months of age. The reduction in swimming velocity was even more pronounced in SCA6 84Q mice which can be attributed to their ataxic symptoms. To visualize the averaged swim patterns of SCA6 84Q mice, 3D heat maps were generated on day 1, 4 and 7 which further illustrate comparable learning behavior of SCA6 84Q mice at all ages tested (Fig. 1E). While on day 1 mice spent approximately equal times in all five potential target alleys, they adapted a goal-directed behavior over the test period of seven days and learned to swim directly to the correct target alley.

Taken together, these results demonstrate that SCA6 84Q mice develop spatial memory deficits in the MWM and the starmaze task which can be interpreted as a sign for cognitive decline. Since the deficits manifest as early as 5 months in the MWM, cognitive decline might even precede motor impairments.

### SCA6 84Q mice exhibit aberrant Purkinje cell simple spike firing

Firing abnormalities of PCs are described to be a common feature for different types of ataxia, including SCA6 (Hoebeek et al., 2005; Walter et al., 2006; Alviña and Khodakhah, 2010; Hansen et al., 2013; Mark et al., 2015). SCA6 84Q mice were also already shown to exhibit reduced firing rates and precision of PCs at 7 months of age when motor deficits start to develop (Jayabal et al., 2016), but little is known about the PC firing properties in earlier stages of the disease. To understand if early-onset spatial memory deficits underlie PCs signaling impairments and to further investigate the development and pathobiology of SCA6, spontaneous PC simple spike firing properties were analyzed in anaesthetized SCA6 84Q and control mice at 3, 5, 7 and 12 months of age (Fig. 3A).

**Figure 3:**
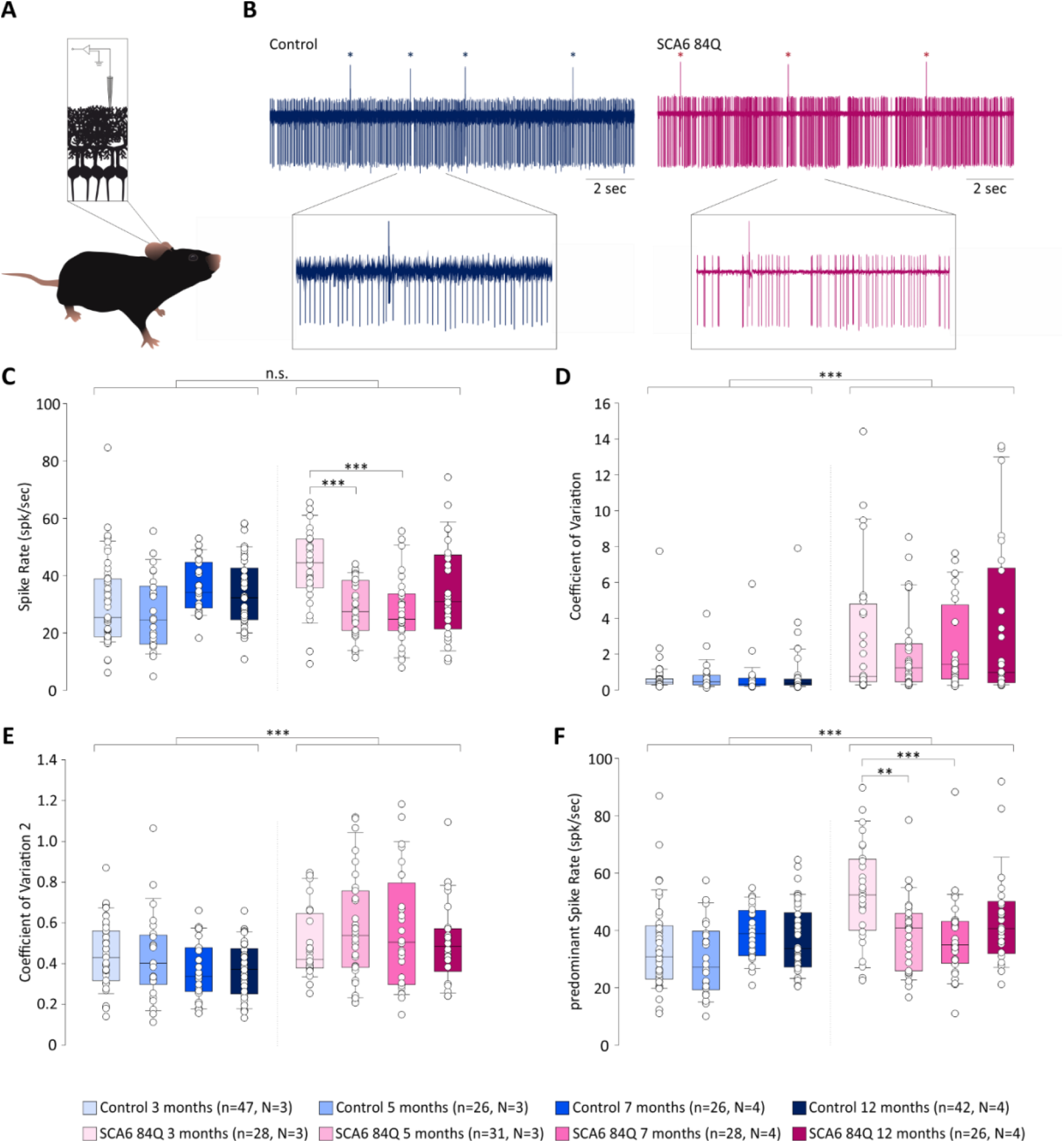
SCA6 84Q mice display aberrant Purkinje cell firing. **(A)** Extracellular recordings of cerebellar PCs were measured in anaesthetized SCA6 84Q and control mice at 3, 5, 7 and 12 months old. **(B)** Example traces from a 12 months old control and SCA6 84Q mouse. SCA6 84Q PCs showed irregular spontaneous simple spike firing. Complex spikes are highlighted by asterisks. PCs of SCA6 84Q and control mice displayed comparable **(C)** mean simple spike firings rates (p=0.355), but exhibited a significant increase in the **(F)** predominant firing rate (p<0.001). Especially the 3 months old SCA6 84Q mice demonstrated augmented mean and predominant simple spike firing frequencies compared to 5 and 7 months old SCA6 84Q mice (p≤0.004). The regularity of PC simple spike firing was diminished in SCA6 84Q mice as shown by an elevated **(D)** coefficient of variation (p<0.001) and **(E)** coefficient of variation 2 (p<0.001). Age-dependent differences were not found for control or SCA6 84Q animals. When applicable data are presented as mean ± SEM. Statistical significance was determined with a two-way ANOVA (*p≤0.05, **p≤0.001, ***p≤0.001). See also extended Table S3.

We found that mean simple spike firing frequencies of PCs were similar between SCA6 84Q (3 months: 42.70 ± 2.59 Hz, 5 months: 28.28 ± 1.73 Hz, 7 months: 27.50 ± 2.32 Hz, 12 months: 34.86 ± 3.32 Hz) and control mice (3 months: 30.68 ± 2.17 Hz, 5 months: 26.73 ± 2.51 Hz, 7 months: 35.78 ± 1.79 Hz, 12 months: 34.00 ± 1.82 Hz, p=0.355, ANOVA, Fig. 3C). Within the two groups, a minor age-dependent increase of simple spike firing was found in control mice (p=0.044, ANOVA), whereas SCA6 84Q mice showed a decrease in simple spike firing frequencies over time (p<0.001, ANOVA). In particular, at 3 months of age simple spike firing rates were significantly elevated compared to 5 and 7 months of age (p<0.001, pairwise comparisons). Next, we evaluated the regularity of firing and found that simple spikes from control PCs fired with high precision (coefficient of variation of interspike intervals (CV) at 3 months: 0.69 ± 0.16, 5 months: 0.73 ± 0.17, 7 months: 0.69 ± 0.22, 12 months: 0.82 ± 0.21, Fig. 3B, D). In contrast SCA6 84Q PCs showed a significantly reduced simple spike firing precision (CV at 3 months: 2.98 ± 0.71, 5 months: 2.07 ± 0.40, 7 months: 2.41 ± 0.46, 12 months: 3.52 ± 0.87, p<0.001, ANOVA, Fig. 3B, D). Since we observed bursting activity in SCA6 84Q mice, we also calculated the coefficient of variation for adjacent intervals of ISIs, CV2, which is a more powerful measure for spike regularity, but found similar results (Fig. 3E). Even during trains of continuous spiking, control PCs exhibited a more regular spiking behavior (3 months: 0.44 ± 0.02, 5 months: 0.43 ± 0.04, 7 months: 0.36 ± 0.03, 12 months: 0.37 ± 0.02) than SCA6 84Q mice (3 months: 0.49 ± 0.03, 5 months: 0.57 ± 0.05, 7 months: 0.57 ± 0.05, 12 months: 0.50 ± 0.04, p<0.001, ANOVA, Fig. 3E). No age-dependent differences in the simple spike firing precision were observed for control (CV: p=0.952, CV2: p=0.062, ANOVA) or SCA6 84Q mice (CV: p=0.375, CV2: p=0.436, ANOVA). Since irregular firing can be reflected in an overall reduction of the mean firing rate caused by spike pauses, we additionally calculated the predominant firing rate (Fig. 3F). In fact, now we discovered increased predominant firing rates in SCA6 84Q mice (3 months: 53.12 ± 3.30, 5 months: 39.05 ± 2.35, 7 months: 36.96 ± 2.72, 12 months: 43.74 ± 3.20) compared to controls (3 months: 33.81 ± 2.17, 5 months: 29.88 ± 2.50, 7 months: 38.88 ± 1.80, 12 months: 37.04 ± 1.83, p<0.001, ANOVA). Similar to the mean firing frequency, SCA6 84Q mice exhibited significantly increased predominant firing frequencies in comparison to older mice at 3 months of age (p≤0.004, pairwise comparisons).

In summary, SCA6 84Q PCs demonstrate a highly irregular spiking behavior at all ages tested that is not accompanied by a decrease of the mean firing frequency. This is achieved by an elevation of the firing rates during trains of continual spiking which presents as an increase of the predominant firing rate.

### SCA6 84Q mice show enhanced swellings of the proximal Purkinje cell axons

Perturbed cell firing is a key sign for neuronal dysfunction which in turn can be associated with morphological changes like the development of axonal swellings (Louis et al., 2014). To determine if the PC firing abnormalities described above are accompanied by morphological changes, the occurrence of axonal swellings in PCs was histologically analyzed at respective ages using calbindin and IP3 receptor antibody staining which was previously described to label the majority of axonal swellings (Lang-Ouellette et al., 2021). Three areas of the cerebellum, Crus I and Crus II hemispheres as well as vermal lobule 3 and 8 were analyzed based on their elevated *cfos* activity during the MWM ((Surdin et al., 2022); Fig. 4A, D, J).

**Figure 4:**
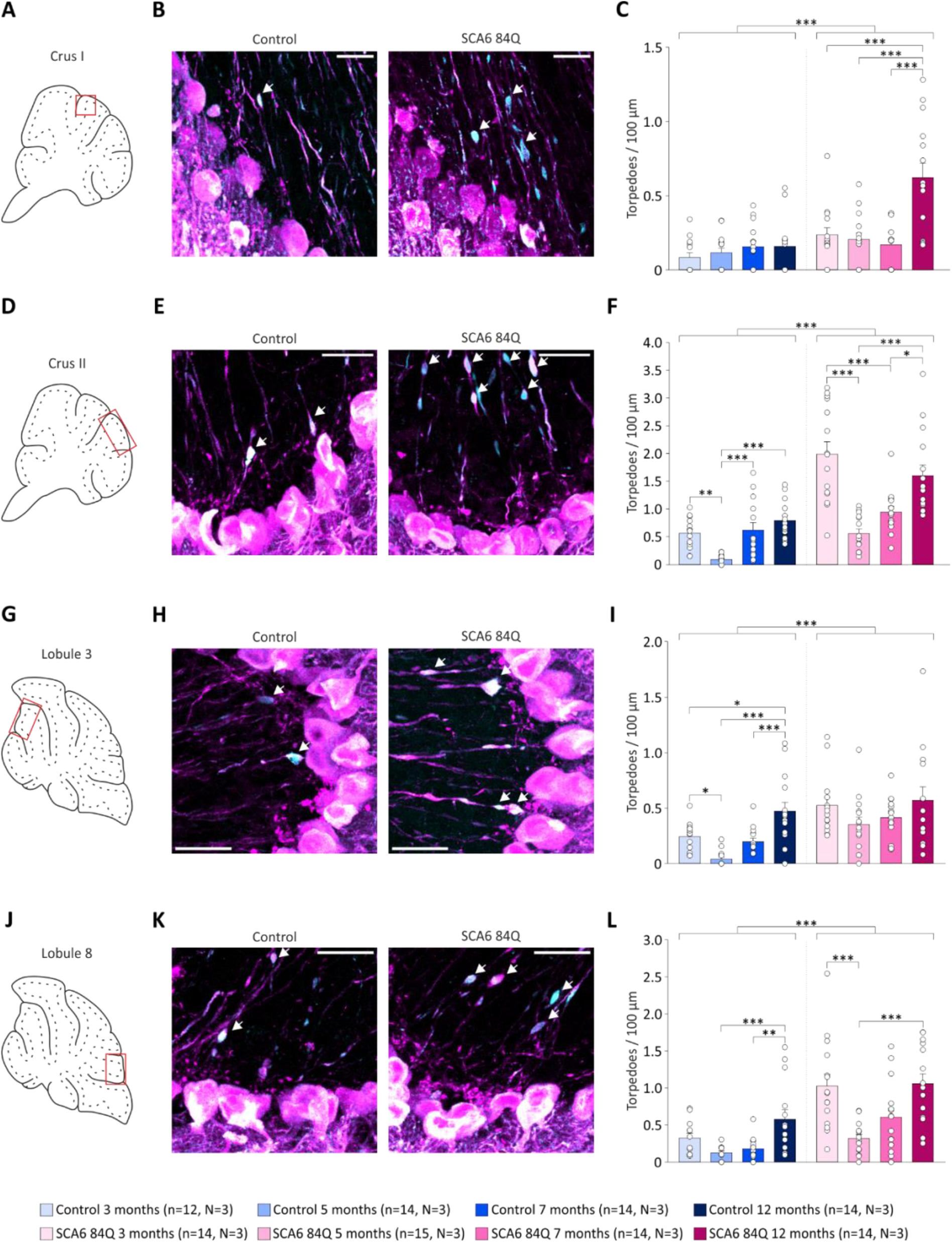
SCA6 84Q mice show increased numbers of axonal swellings. **(A, D, G, J)** Scheme of cerebellar sections highlighting (red boxes) the analyzed areas in Crus I, Crus II, lobule 3 and lobule 8 of axonal swellings. For each area 5 images/brain were analyzed for the number of axonal swellings/100 µm. Example images of PCs (magenta, labeled with calbindin) illustrate more torpedoes (cyan, labeled with IP3 receptor) in **(B)** Crus I, **(E)** Crus II, **(H)** lobule 3 and **(K)** lobule 8 for SCA6 84Q compared to control mice. Scale bar indicates 30 µm. Axonal swellings are marked with arrows. Quantification confirmed that SCA6 84Q mice exhibited more torpedoes compared to controls in **(C)** Crus I (p<0.001) and **(F)** Crus II (p<0.001) of the cerebellar hemispheres as well as in **(I)** lobule 3 (p<0.001) and **(L)** lobule 8 (p<0.001) of the vermis. For both SCA6 84Q and control mice elevated numbers of torpedoes were found at 12 month of age for various brains regions. SCA6 84Q mice additionally displayed an increa sed torpedo occurrence at 3 month of age in Crus II and lobule 8. When applicable data are presented as mean ± SEM. Number of individual slices analyzed are shown below the bar graphs. Statistical significance was determined with a two-way ANOVA (*p≤0.05, **p≤0.001, ***p≤0.001). See also extended Table S4.

We found a significantly higher torpedo occurrence in the anterior Crus I of SCA6 84Q mice (3 months: 0.24 ± 0.06, 5 months: 0.21 ± 0.04, 7 months: 0.16 ± 0.05, 12 months: 0.62 ± 0.14) than in controls (3 months: 0.1 ± 0.04, 5 months: 0.2 ± 0.04, 7 months: 0.16 ± 0.02, 12 months: 0.16 ± 0.02, p<0.001, ANOVA, Fig. 4B, C). No age-dependent differences in torpedo occurrence was observed in the control group (p=0.489, ANOVA), while the number of axonal swellings differed in SCA6 84Q mice among the ages (p<0.001, ANOVA). Especially, 12-months-old SCA6 84Q mice exhibited more axonal swelling in comparison to younger SCA6 84Q mice (p<0.001, ANOVA). Axonal swellings in the posterior Crus II of SCA6 84Q mice (3 months: 2.00 ± 0.51, 5 months: 0.56 ± 0.09, 7 months: 0.97 ± 0.18, 12 months: 1.60 ± 0.43) were also elevated compared to controls (3 months: 0.56 ± 0.13, 5 months: 0.10 ± 0.01, 7 months: 0.59 ± 0.32, 12 months: 0.81 ± 0.14, p<0.001, ANOVA, Fig. 4E, F). Contrary to Crus I, the posterior Crus II region displayed diverging torpedo occurrences across the different ages of both genotypes (p<0.001 for both genotypes, ANOVA). While in the control group especially 5 months old mice displayed low numbers of axonal swellings (p≤0.004, pairwise comparison), SCA6 84Q mice showed a twofold increase of torpedoes at 3 and 12 months of age (p≤0.045, pairwise comparison).

Similar results were found in the cerebellar vermis. The anterior lobule 3 area also showed an augmented number of torpedoes in SCA6 84Q (3 months: 0.52 ± 0.09, 5 months: 0.35 ± 0.12, 7 months: 0.41 ± 0.04, 12 months: 0.60 ± 0.24) mice compared to controls (3 months: 0.23 ± 0.05, 5 months: 0.04 ± 0.02, 7 months: 0.20 ± 0.05, 12 months: 0.47 ± 0.17, p<0.001, ANOVA, Fig. 4H, I). However, unlike the results from the hemispheres, we found age-dependent differences for control but not SCA6 84Q mice (Control: p<0.001, SCA6: p=0.212, ANOVA) from lobule 3. Moreover, a notable increase of axonal swellings was found in 12 months old control mice (p≤0.012, pairwise comparison). In agreement with the results from CrusI/II and lobule 3, an accumulation of torpedoes in SCA6 mice (3 months: 1.03 ± 0.26, 5 months: 0.32 ± 0.07, 7 months: 0.61 ± 0.23, 12 months: 1.06 ± 0.30) compared to the control group (3 months: 0.32 ± 0.10, 5 months: 0.13 ± 0.02, 7 months: 0.18 ± 0.05, 12 months: 0.58 ± 0.24, p<0.001, ANOVA, Fig. 4K, L) was noted in lobule 8. Again, we found diverging torpedo occurrences across the different ages of both genotypes in lobule 8 (p<0.001 for both genotypes, ANOVA). In agreement with the results for vermal lobule 3, control mice exhibited elevated numbers of torpedoes at 12 months of age (p≤0.003, pairwise comparison). On the other hand, the torpedo occurrence in lobule 8 of SCA6 84Q mice was increased in both 3 and 12 months old mice (p<0.001).

Together, the data indicate that SCA6 84Q PCs in addition to cell firing failures also exhibit morphological abnormalities in terms of an increased torpedo occurrence. While an increase in the number of axonal swellings could be observed for both control and SCA6 84Q mice at 12 month of age in various regions of the cerebellum, SCA6 84Q mice additionally showed an increase of torpedoes at 3 months of age in the posterior lobule 8 and Crus II.

### Gq-protein stimulation of the cerebellum rescues spatial navigation impairments in SCA6 84Q mice

PKC-dependent LTD at the PF-PC synapse has been previously shown to be involved in spatial memory processes that heavily depend on the procedural component of navigation using the L7-PKCI mouse model (Burguière et al., 2005). Impaired cerebellar synaptic plasticity including LTD was previously found in another SCA6 mouse model and proposed to be part of the SCA6 disease pathobiology as well (Mark et al., 2015). Since cognitive and motor learning in the cerebellum is most likely controlled by LTD and LTD is mediated by the mGluR1 dependent Gq-protein signaling (Hartmann et al., 2004), we aimed to improve the goal-directed navigation of SCA6 84Q mice by stimulating the cerebellar Gq pathway with DREADDs (Fig. 5A, D).

**Figure 5:**
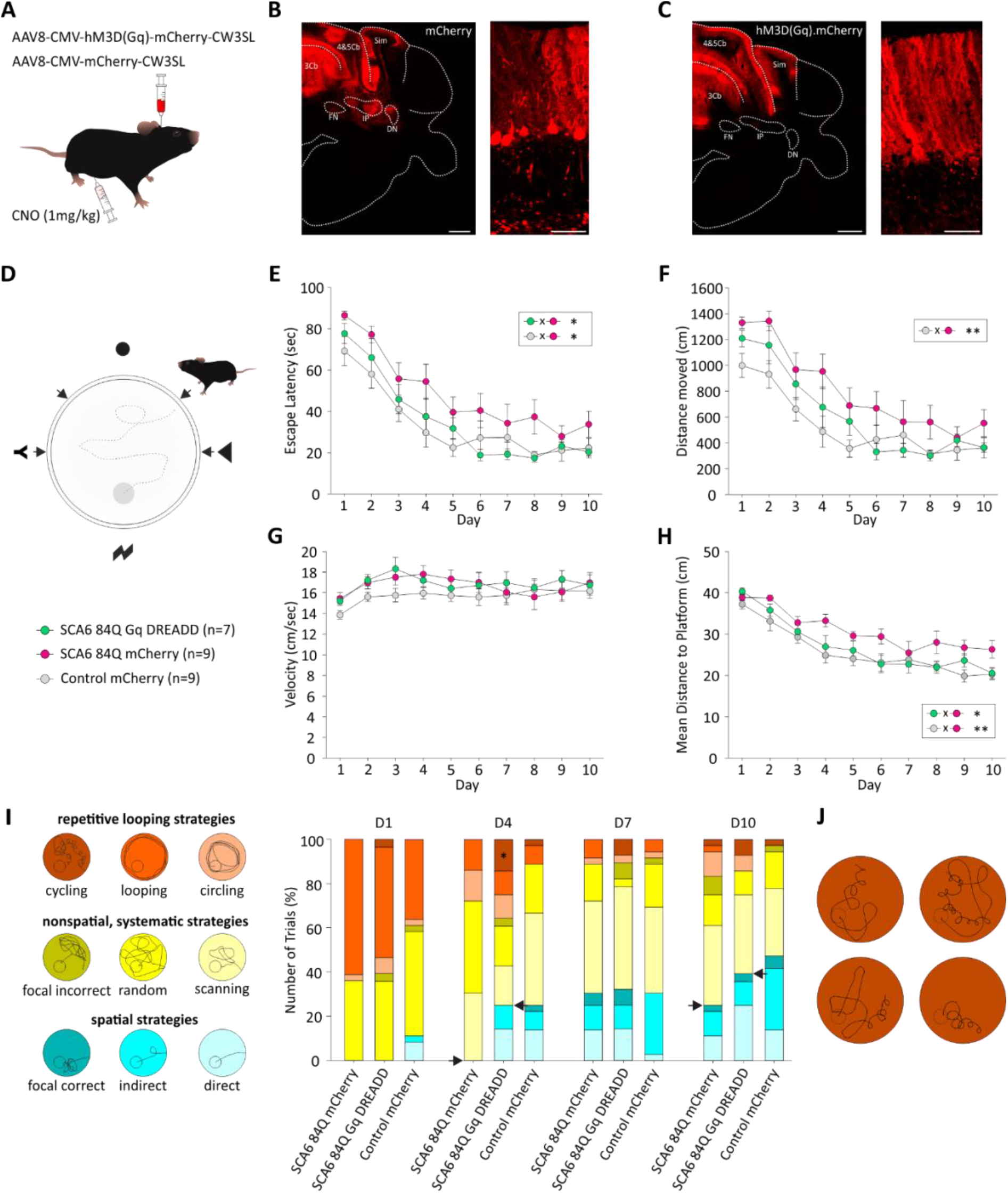
Rescue of spatial navigation impairments in SCA6 84Q mice by enhancing cerebellar Gq-protein signaling. **(A)** SCA6 84Q mice were injected with either AAV8-CMV-hM3D(Gq)-mCherry-CW3SL or AAV8-mCherry-CW3SL as a control. Control mice were injected with AAV8-mCherry-CW3SL only. Thirty min prior to the experiment CNO (1 mg/kg) was administered intraperitoneally. Example images of **(B)** mCherry and hM3D(Gq)-mCherry **(C)** expression in predominantly cerebellar PCs. Scale bars indicate 500 µm in the overview and 50 µm in the close-up. **(D)** Schematic overview of the MWM. Mice searched for a hidden platform in an aquatic open field using external visual cues (circle, triangle, lightning, Y) located at the 4 cardinal directions. Mice were trained for 10 consecutive days consisting of 4 trials/day from 4 pseudorandomized starting positions (marked by arrows). All 3 groups, SCA6 84Q mCherry, SCA6 84Q Gq-DREADD and control mCherry mice, successfully learned to find the platform over the 10 day training period indicated by a continuous decrease in the **(E)** escape latency, **(F)** overall distance moved and **(H)** mean distance to the platform (p<0.001 for all). Gq-DREADD activation was able to improve spatial learning of SCA6 84Q mice (SCA6 84Q Gq-DREADD) to control (control mCherry) levels as both groups did not differ in the above mentioned parameters (p>0.05). Furthermore, significant improvements compared to SCA6 84Q mCherry mice were found for the escape latency (p=0.049) and mean distance to the platform (p=0.048). **(G)** No difference in the swimming speeds was found between groups (p=0.413) indicating normal motor behavior. **(I)** Evaluation of the navigation strategies from SCA6 84Q and control swimming paths on days 1, 4, 7 and 10. Each trial was assigned to 1 of 9 different search strategies classified into 3 superior categories: spatial strategies, systematic but nonspatial strategies and repetitive looping strategies. In general, over the 10 day test period successful learning of the task was illustrated by a decrease of repetitive looping strategies and an increase of spatial strategies. Gq-DREADD dependent recovery of strategic learning was most evident on day 4 and 10 (marked with arrows), where SCA6 84Q Gq-DREADD mice used more spatial and less nonstrategic (nonspatial and looping) strategies compared to SCA6 84Q mCherry mice. **(J)** SCA6 84Q Gq-DREADD mice adapted a goal-directed circling behavior that was not seen in the other groups and makes up 75 % of the cycling behavior on day 4 (marked with * in (I)). When applicable, data are presented as mean ± SEM. Statistical significance was determined with a two-way ANOVA with repeated measures (*p≤0.05, **p≤0.001, ***p≤0.001). See also extended Table S5. 4&5Cb, lobule 4&5; 3Cb, lobule 3; Sim, simple lobule; FN, fastigial nucleus; IP, interposed nucleus; DN, dentate nucleus.

Both the Gq-DREADD (hM3D(Gq)-mCherry) and a control mCherry were robustly expressed in the anterior cerebellar cortex of 5 month old mice, predominantly in PCs of lobules 3, 4 and 5 and the simple lobule (Figure 5B, C). Expression of the viral constructs did not interfere with spatial learning in the MWM because all 3 groups (control mCherry, SCA6 84Q mCherry, SCA6 84Q Gq-DREADD) successfully learned to find the platform over the 10 day test period indicated by a continuous decrease in the time (p<0.001 for all 3 groups, ANOVA, Fig. 5E), overall distance moved (p<0.001 for all 3 groups, ANOVA, Fig. 5F) and mean distance (p<0.001 for all 3 groups, ANOVA, Fig. 5H) to the hidden platform. As expected, SCA6 84Q mCherry mice performed poorly in all 3 parameters (p≤0.017, pairwise comparisons) compared to control mCherry mice, confirming the spatial navigation impairments of SCA6 84Q mice. Cerebellar stimulation of Gq with DREADDs significantly improved the spatial navigation performance of SCA6 84Q mice to control levels. The improvements predominantly manifested in a lower escape latency (p=0.049, pairwise comparison) and mean distance to the platform (p=0.048, pairwise comparison) compared to SCA6 84Q mCherry mice. The swimming velocity, however, did not differ between the 3 groups (p=0.413, ANOVA, Fig. 5G) indicating that the Gq-DREADD stimulation did not interfere with the general motor performance. Individual swimming paths were analyzed to determine their searching strategies (Fig. 5I). Overall, successful learning of the MWM task was demonstrated by a decrease in repetitive looping strategies and an increase in spatial strategies during the 10 day testing period. While comparable searching strategies for SCA6 84Q mCherry and SCA6 84Q Gq-DREADD were found on day 1, Gq-DREADD dependent improvement of SCA6 84Q Gq-DREADD mice was most prominent on day 4 and 10. On day 4 SCA6 84Q mCherry mice did not show any spatial strategies and predominantly engage in nonspatial, systematic strategies, whereas SCA6 84Q Gq-DREADD and control mCherry mice showed similar numbers of spatial strategies. On day 10 SCA6 84Q Gq-DREADD mice displayed more spatial and less nonspatial strategies compared to SCA6 84Q mCherry mice in the MWM. Interestingly, upon Gq-DREADD stimulation SCA6 84Q mice developed a goal-directed circling behavior which was not observed in other mouse lines and makes up 75 % of the cycling behavior seen on day 4 (Fig. 5J).

Taken together, these results suggest that spatial navigation impairments of SCA6 84Q mice are caused by a reduced cerebellar Gq-protein signaling and thus potentially reduced cerebellar LTD, supporting the idea that disruption of synaptic plasticity is a key feature of the SCA6 pathobiology.

## Discussion

In the present study we show that SCA6 84Q mice exhibit spatial navigation deficits in the MWM and starmaze. While deficits were mild in the starmaze, we found strong impairments in the MWM starting at 5 months of age. Furthermore, we rescued spatial memory deficits by enhancing Gq-protein signaling in the cerebellum, indicating a potential disruption of Gq-protein signaling and synaptic plasticity in SCA6 84Q mice. Increased irregularity of PC firing and axonal swellings in SCA6 84Q PCs further confirm a disruption in PC function which was observed as early as 3 months of age and largely independent of disease progression.

It is still an ongoing debate to what extend SCA6 patients suffer from cognitive deficiencies. Mild cognitive dysfunctions which include impairments in visual-motor processing, verbal fluency, executive functions and social cognition have been described so far but the body of evidence is continuously growing (Suenaga et al., 2008; Pereira et al., 2017; Giocondo and Curcio, 2018; Abdelgabar et al., 2019). Recently, a decrease in innate fear and defense behavior has for the first time been described in a SCA6 mouse model (Bohne et al., 2022). In this study we were able demonstrate that cognitive decline, in particular spatial navigation deficits, is in fact a feature of the SCA6 pathology. Spatial navigation is a cognitive task that requires the integration of self-motion and external sensori-motor information to build an internal map of the surrounding context (declarative component) which is then used to optimize the goal-directed path (procedural component). The cerebellum is supposed to contribute to this process providing self-motion information and optimizing goal-directed navigation (Rochefort et al., 2013). Accordingly, it was shown that PC degeneration leads to deficits in spatial memory formation (Goodlett et al., 1992; Martin et al., 2003) which corroborates the spatial navigation deficits we showed in SCA6 84Q mice. The spatial navigation impairments in 3 - 7 month old SCA6 84Q mice cannot be attributed to motor deficits, since SCA6 84Q mice develop ataxia at the age of 7 months (Watase et al., 2008; Jayabal et al., 2015) and displayed comparable or even enhanced swimming velocities in the MWM (Fig 1D). Similarly, these mice only develop a minor swimming deficit in their hindlimb kicks which did not alter their swimming speed (Jayabal et al., 2015). The difference in the age of onset for motor and cognitive symptoms can be caused by different vulnerabilities of underlying cerebellar microcircuits to the SCA6 pathology. Such a pattern was for example found in autosomal-recessive spastic ataxia of Charlevoix-Saguenay mice which display an enhanced PC death especially in zebrin-negative PCs in anterior lobules. These data suggest that the vulnerability of PCs to degeneration relies on the location and molecular identity (Toscano Márquez et al., 2021). While motor functions are predominantly processed in vermal lobules IV-VI, cognition and in particular spatial processing involves the Crus I/II hemispheres (Kelly and Strick, 2003; Stoodley et al., 2012; Iglói et al., 2015; Watson et al., 2019). In the MWM increased PC activity was found in the entire hemispheres as well as vermal lobules 3-5 and 8/9 (Surdin et al., 2022). However, in SCA6 patients PC loss and cerebellar atrophy is described to be most prominent in the vermis and less prominent in the hemispheres (Satoh et al., 1998; Lukas et al., 2006). Surprisingly, we found strong spatial memory deficits in the MWM but only mild deficits in the starmaze. Similar results were found for L7-PKCI mice which lack LTD at the PF-PC synapse (Zeeuw et al., 1998; Burguière et al., 2005). This discrepancy is attributed to the distinct characteristics of the tasks. Both tasks require the declarative component of spatial memory while the procedural component is only required in the MWM suggesting that L7-PKCI mice cannot adapt their goal-directed behavior and that LTD at the PF-PC is necessary for this process. Since LTD dysfunction at the PF-PC synapse was previously reported for another SCA6 mouse line (Mark et al., 2015), the deficits we found for SCA6 84Q mice may also be caused by disruptions in LTD. Rescue of spatial memory deficits via cerebellar Gq-protein stimulation, a key pathway for LTD induction at the PF-PC synapse (Hartmann et al., 2004), indirectly confirmed that synaptic plasticity is impaired in SCA6 84Q mice.

For numerous spinocerebellar ataxias including SCA6 the development of motor deficits is strongly associated with abnormal PC firing. While reduced firing rates were found in SCA1 (Hourez et al., 2011) and SCA2 (Hansen et al., 2013) mice, PC firing deficits were observed as reduced firing precision (Mark et al., 2015) or as a combination of both reduced firing rate and precision (Jayabal et al., 2016) in different mouse models for SCA6. Consistent with the observations made by (Jayabal et al., 2016), we also report aberrant PC simple spike firing precision in SCA6 84Q mice at 7 months of age. In contrast to their study, we did not find any reduced firing frequencies. More importantly, we show that PC firing precision deficits already appear in an early, presymptomatic disease state as early as 3 months, suggesting that they do not ultimately induce motor symptoms or cognitive decline which manifest at 7- and 5- months of age respectively. However, we cannot exclude the possibility that there might be early fine motor and cognitive deficiencies which we were not able to detect with our behavioral tests. Presymptomatic changes in cell signaling have been described for other disease such as SCA2 (Hansen et al., 2013) and Huntington’s disease (Dougherty et al., 2012). In both cases a decrease in PC firing frequencies has been shown that precedes the development of behavioral and morphological abnormalities and is supposed to be a mild initial stage which impact becomes worse with aging. Nevertheless, the PC firing deficits found by us and others do not reflect channel dysfunction per se. Since the same voltage sensitivity as well as activation and deactivation kinetics were described for the P/Q-type Ca^2+^-channel in SCA6 84Q mice (Watase et al., 2008). A more promising explanation for the simple spike firing deficits is that the P/Q-type Ca^2+^-channel exerts a dominant-negative effect on interaction partners. Especially, an altered coupling to Ca^2+^-activated K^+^-channels which form complexes with P/Q-type Ca^2+^-channels (Berkefeld et al., 2006) and mediate the fast hyperpolarization thereby enabling high-frequency repetitive firing results in altered firing precision (Womack and Khodakhah, 2002, 2003; Walter et al., 2006). Correspondingly, the irregular firing behavior that was found in another SCA6 mouse line can be mimicked via application of Ca^2+^-activated K^+^-channel blockers in control PCs (Mark et al., 2015). Additionally, it was shown that chlorzoxazone, an activator of Ca^2+^-activated K^+^-channels, restores PC firing deficits and reduces motor symptoms in another P/Q-type Ca^2+^-channel mouse mutant (Alviña and Khodakhah, 2010), and blockade of voltage-dependent K^+^-channels using 4-aminopyridine successfully enhanced PC firing precision and partially rescues ataxia in SCA6 84Q mice (Jayabal et al., 2016).

Besides being a normal feature of cerebellar maturation (Gravel et al., 1986; Ljungberg et al., 2016) the formation of axonal swellings in PCs is commonly found in neurodegenerative disease like essential tremor, SCAs or even multiple sclerosis (Suzuki et al., 1969; Louis et al., 2014). For SCA6 84Q mice a normal development of torpedoes was described during maturation of the cerebellum at P11 while increased numbers of axonal swellings were reported in lobule 3 after 2 years (Ljungberg et al., 2016). Since PC loss was also found in 2-year-old SCA6 84Q mice these result lead to the suggestion, that the development of torpedoes contributes to the neurodegeneration process as it was previously predicted (Manor et al., 1991; Maia et al., 2015). In contrast to these results, we were able to demonstrate elevated numbers of axonal swellings in SCA6 84Q mice at earlier ages. Similar to our electrophysiological results, we detected elevated torpedo counts also as early as 3 months of age, raising the idea that the development of axonal swellings is closely linked to aberrant PC firing. In fact, it was recently shown that PC axonal swellings enhance the fidelity of actions potentials and improve cerebellar function (Lang-Ouellette et al., 2021). Even though these experiments were conducted in young, healthy animals, there are supposedly subtle structural differences between developmental and disease-related torpedoes in PCs (Ljungberg et al., 2016). It can be hypothesized that rather than playing a pathophysiological role, axonal swellings develop as a compensatory mechanism during cellular stress (Mann et al., 1980). In conclusion, we were able to demonstrate a direct link between cognitive decline and the SCA6 pathobiology which is the result of deficient PC signaling and presumably synaptic plasticity. Our findings need to be confirmed in SCA6 patients, but we provide the first evidence in mice that cognitive decline might even develop earlier than motor deficits for SCA6. This raises awareness to survey cognitive abnormalities during clinical examination of SCA6 patients to potentially diagnose and prevent the progression of cognitive deficiencies in earlier disease stages and to optimize the individual treatment strategy by enhancing PC activity. This is of special interest because in contrast to mouse models, symptoms appear at a broad range of age in SCA6 patients (Gomez et al., 1997).

## Author contributions

MDM and MG designed project and experiments; MG, HS, CDCT and LN performed experiments; MG analyzed data; MDM and MG wrote the paper.

## Ethics approval

Maintenance and experimental procedures of animals were conducted with approval of a local ethics committee (Bezirksamt Arnsberg) and the animal care committee of North Rhine-Westphalia, Germany, based at the LANUV (LANUV; Landesamt für Umweltschutz, Naturschutz und Verbraucherschutz Nordrhein-Westfalen, Germany). The study was carried out in accordance with the European Communities Council Directive of 2010 (2010/63/ EU) for care of laboratory animals and supervised by the animal welfare commission of the Ruhr-University Bochum.

## Data Availability Statement

All data supporting this study are available from the corresponding author upon request.

## Acknowledgements

We would like to thank Margareta Möllmann, Stephanie Krämer, Petra Knipschild, Wolfgang Kruse, Winfried Junke, Kevin Sowa, Stefan Dobers, Gina Hillgruber, Nicole Ozdowski and Manuela Schmidt for their excellent technical assistance. This work was supported by the Deutsche Forschungsgemeinschaft (DFG; German Funding Foundation) MA 5806/2-1 (MDM), Project number 316803389-SFB1280 (Project A21, MDM) and MA 5806/7-1. MG was supported by MA 5806/2-1 (MDM).

## Extended Data

**Table S1.** Statistics for the performance of SCA6 84Q and control mice in the Morris water maze: repeated measures two-way ANOVA.

| Factor | df<sub>num</sub> | df<sub>den</sub> | F | p |
| --- | --- | --- | --- | --- |
| **Escape Latency** |  |  |  |  |
| Genotype | 1 | 83 | 13.608 | <0.001 |
| Time (SCA6) | 6.009<sup>1</sup> | 228.333<sup>1</sup> | 49.947<sup>1</sup> | <0.001<sup>1</sup> |
| Time (Control) | 5.957<sup>1</sup> | 268.049<sup>1</sup> | 65.296<sup>1</sup> | <0.001<sup>1</sup> |
| Age (SCA6) | 3 | 38 | 15.815 | <0.001 |
| Age (Control) | 3 | 45 | 1.570 | 0.21 |
| Pairwise comparisons (p≤0.05 only): |  |  |  |  |
| 3 months x 5 months (SCA6) |  |  |  | 0.002 |
| 3 months x 7 months (SCA6) |  |  |  | 0.049 |
| 3 months x 12 months (SCA6) |  |  |  | <0.001 |
| 7 months x 12 months (SCA6) |  |  |  | 0.002 |
| **Distance moved** |  |  |  |  |
| Genotype | 1 | 83 | 20.252 | <0.001 |
| Time (SCA6) | 5.964<sup>1</sup> | 226.629<sup>1</sup> | 48.827<sup>1</sup> | <0.001<sup>1</sup> |
| Time (Control) | 6.301<sup>1</sup> | 283.536<sup>1</sup> | 66.779<sup>1</sup> | <0.001<sup>1</sup> |
| Age (SCA6) | 3 | 38 | 5.265 | 0.004 |
| Age (Control) | 3 | 45 | 3.461 | 0.024 |
| Pairwise comparisons (p≤0.05 only): |  |  |  |  |
| 3 months x 5 months (SCA6) |  |  |  | 0.01 |
| 3 months x 7 months (SCA6) |  |  |  | 0.04 |
| 3 months x 12 months (SCA6) |  |  |  | 0.027 |
| 3 months x 7 months (Control) |  |  |  | 0.048 |
| **Velocity** |  |  |  |  |
| Genotype | 1 | 83 | 7.185 | 0.009 |
| Time (SCA6) | 5.316<sup>1</sup> | 201.990<sup>1</sup> | 1.463<sup>1</sup> | 0.2<sup>1</sup> |
| Time (Control) | 5.964<sup>1</sup> | 268.378<sup>1</sup> | 1.000<sup>1</sup> | 0.135<sup>1</sup> |
| Age (SCA6) | 3 | 38 | 9.746 | <0.001 |
| Age (Control) | 3 | 45 | 5.912 | 0.002 |
| Pairwise comparisons (p≤0.05 only): |  |  |  |  |
| 3 months x 12 months (SCA6) |  |  |  | <0.001 |
| 5 months x 12 months (SCA6) |  |  |  | 0.009 |
| 7 months x 12 months (SCA6) |  |  |  | <0.001 |
| 3 months x 5 months (Control) |  |  |  | 0.002 |
| 3 months x 7 months (Control) |  |  |  | 0.021 |
| 3 months x 12 months (Control) |  |  |  | 0.015 |

<sup>1</sup>Greenhouse-Geisser correction

**Table S2.** Statistics for the performance of SCA6 84Q and control mice in the starmaze: repeated measures two-way ANOVA.

| Factor | df<sub>num</sub> | df<sub>den</sub> | F | p |
| --- | --- | --- | --- | --- |
| **Escape Latency** |  |  |  |  |
| Genotype | 1 | 62 | 4.627 | 0.035 |
| Time (SCA6) | 4.184<sup>1</sup> | 129.708<sup>1</sup> | 11.434<sup>1</sup> | <0.001<sup>1</sup> |
| Time (Control) | 3.588<sup>1</sup> | 111.225<sup>1</sup> | 15.731<sup>1</sup> | <0.001<sup>1</sup> |
| Age (SCA6) | 3 | 31 | 3.651 | 0.023 |
| Age (Control) | 3 | 31 | 5.810 | 0.003 |
| Pairwise comparisons (p≤0.05 only): |  |  |  |  |
| 7 months x 12 months (SCA6) |  |  |  | 0.039 |
| 3 months x 12 months (Control) |  |  |  | 0.003 |
| 5 months x 12 months (Control) |  |  |  | 0.029 |
| **Distance moved** |  |  |  |  |
| Genotype | 1 | 62 | 6.732 | 0.012 |
| Time (SCA6) | 4.146<sup>1</sup> | 128.529<sup>1</sup> | 6.851<sup>1</sup> | <0.001<sup>1</sup> |
| Time (Control) | 4.265<sup>1</sup> | 132.203<sup>1</sup> | 9.502<sup>1</sup> | <0.001<sup>1</sup> |
| Age (SCA6) | 3 | 31 | 3.091 | 0.041 |
| Age (Control) | 3 | 31 | 2.889 | 0.051 |
| Pairwise comparisons (p≤0.05 only): |  |  |  |  |
| 3 months x 7 months (SCA6) |  |  |  | 0.038 |
| **Velocity** |  |  |  |  |
| Genotype | 1 | 62 | 0.346 | 0.559 |
| Time (SCA6) | 3.839<sup>1</sup> | 119.020<sup>1</sup> | 9.064<sup>1</sup> | <0.001<sup>1</sup> |
| Time (Control) | 3.386<sup>1</sup> | 104.957<sup>1</sup> | 24.719<sup>1</sup> | <0.001<sup>1</sup> |
| Age (SCA6) | 3 | 31 | 32.231 | <0.001 |
| Age (Control) | 3 | 31 | 6.900 | 0.001 |
| Pairwise comparisons (p≤0.05 only): |  |  |  |  |
| 3 months x 12 months (SCA6) |  |  |  | <0.001 |
| 5 months x 12 months (SCA6) |  |  |  | <0.001 |
| 7 months x 12 months (SCA6) |  |  |  | <0.001 |
| 3 months x 12 months (Control) |  |  |  | 0.004 |
| 5 months x 12 months (Control) |  |  |  | 0.003 |
| 7 months x 12 months (Control) |  |  |  | 0.039 |

<sup>1</sup>Greenhouse-Geisser correction

**Table S3.**
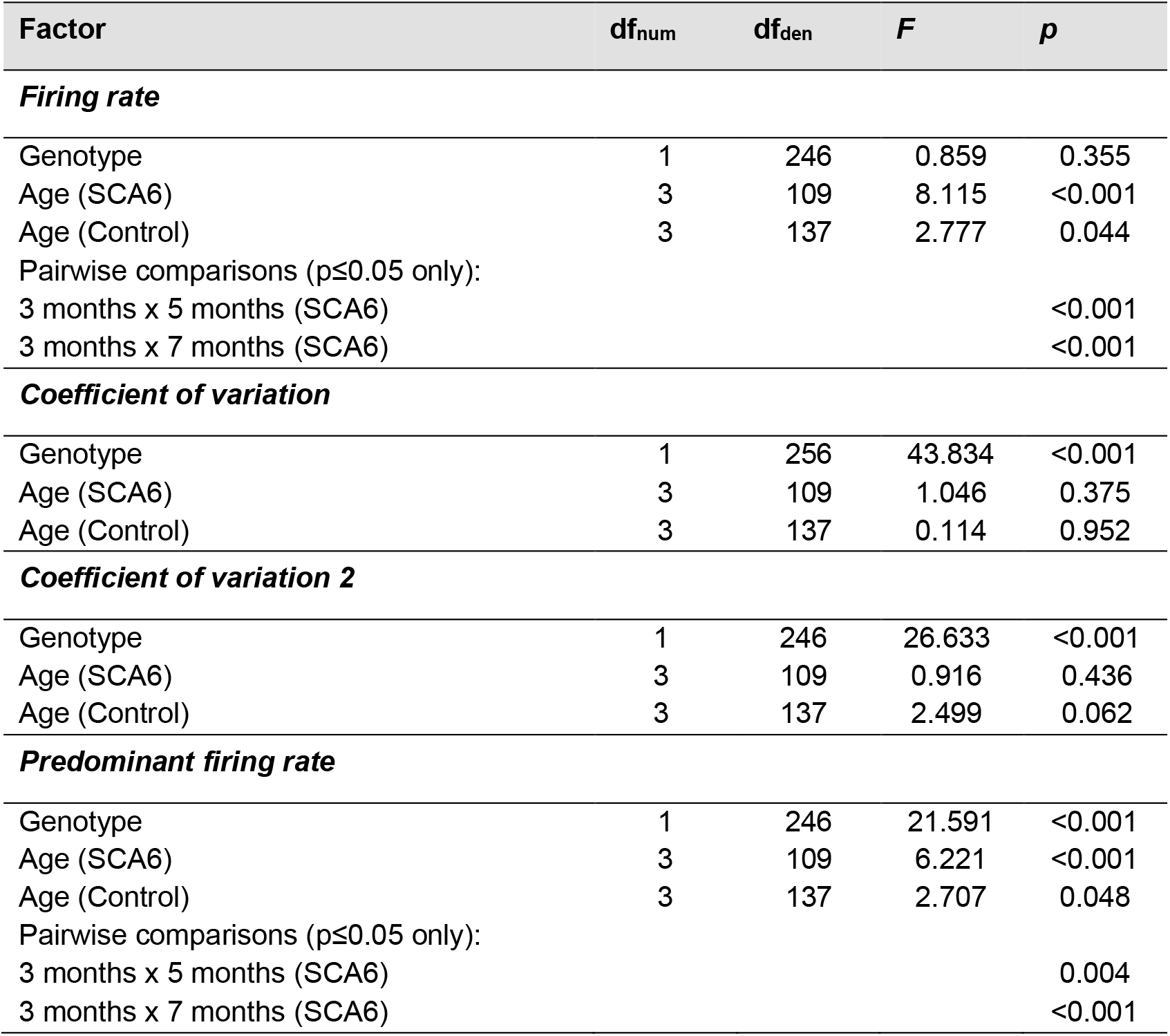
Statistics for the electrophysiological properties of Purkinje cells in SCA6 84Q and control mice: two-way ANOVA.

| Factor | df<sub>num</sub> | df<sub>den</sub> | F | p |
| --- | --- | --- | --- | --- |
| ***Firing rate*** |  |  |  |  |
| Genotype | 1 | 246 | 0.859 | 0.355 |
| Age (SCA6) | 3 | 109 | 8.115 | <0.001 |
| Age (Control) | 3 | 137 | 2.777 | 0.044 |
| Pairwise comparisons (p≤0.05 only): |  |  |  |  |
| 3 months x 5 months (SCA6) |  |  |  | <0.001 |
| 3 months x 7 months (SCA6) |  |  |  | <0.001 |
| ***Coefficient of variation*** |  |  |  |  |
| Genotype | 1 | 256 | 43.834 | <0.001 |
| Age (SCA6) | 3 | 109 | 1.046 | 0.375 |
| Age (Control) | 3 | 137 | 0.114 | 0.952 |
| ***Coefficient of variation 2*** |  |  |  |  |
| Genotype | 1 | 246 | 26.633 | <0.001 |
| Age (SCA6) | 3 | 109 | 0.916 | 0.436 |
| Age (Control) | 3 | 137 | 2.499 | 0.062 |
| ***Predominant firing rate*** |  |  |  |  |
| Genotype | 1 | 246 | 21.591 | <0.001 |
| Age (SCA6) | 3 | 109 | 6.221 | <0.001 |
| Age (Control) | 3 | 137 | 2.707 | 0.048 |
| Pairwise comparisons (p≤0.05 only): |  |  |  |  |
| 3 months x 5 months (SCA6) |  |  |  | 0.004 |
| 3 months x 7 months (SCA6) |  |  |  | <0.001 |

**Table S4.** Statistics for the axonal swellings count in Purkinje cells of SCA6 84Q and control mice: two-way ANOVA.

| Factor | df<sub>num</sub> | df<sub>den</sub> | F | p |
| --- | --- | --- | --- | --- |
| ***Crus I*** |  |  |  |  |
| Genotype | 1 | 107 | 23.580 | <0.001 |
| Age (SCA6) | 3 | 55 | 11.727 | <0.001 |
| Age (Control) | 3 | 53 | 0.819 | 0.489 |
| Pairwise comparisons (p≤0.05 only): |  |  |  |  |
| 3 months x 12 months (SCA6) |  |  |  | <0.001 |
| 5 months x 12 months (SCA6) |  |  |  | <0.001 |
| 7 months x 12 months (SCA6) |  |  |  | <0.001 |
| ***Crus II*** |  |  |  |  |
| Genotype | 1 | 107 | 62.327 | <0.001 |
| Age (SCA6) | 3 | 55 | 15.340 | <0.001 |
| Age (Control) | 3 | 52 | 11.124 | <0.001 |
| Pairwise comparisons (p≤0.05 only): |  |  |  |  |
| 3 months x 5 months (SCA6) |  |  |  | <0.001 |
| 3 months x 7 months (SCA6) |  |  |  | <0.001 |
| 5 months x 12 months (SCA6) |  |  |  | <0.001 |
| 7 months x 12 months (SCA6) |  |  |  | 0.045 |
| 3 months x 5 months (Control) |  |  |  | 0.004 |
| 5 months x 7 months (Control) |  |  |  | <0.001 |
| 5 months x 12 months (Control) |  |  |  | <0.001 |
| ***Lobule 3*** |  |  |  |  |
| Genotype | 1 | 107 | 23.317 | <0.001 |
| Age (SCA6) | 3 | 54 | 1.552 | 0.212 |
| Age (Control) | 3 | 53 | 14.255 | <0.001 |
| Pairwise comparisons (p≤0.05 only): |  |  |  |  |
| 3 months x 5 months (Control) |  |  |  | 0.039 |
| 3 months x 12 months (Control) |  |  |  | 0.012 |
| 5 months x 12 months (Control) |  |  |  | <0.001 |
| 7 months x 12 months (Control) |  |  |  | <0.001 |
| ***Lobule 8*** |  |  |  |  |
| Genotype | 1 | 107 | 36.943 | <0.001 |
| Age (SCA6) | 3 | 56 | 8.389 | <0.001 |
| Age (Control) | 3 | 51 | 6.999 | <0.001 |
| Pairwise comparisons (p≤0.05 only): |  |  |  |  |
| 3 months x 5 months (SCA6) |  |  |  | <0.001 |
| 5 months x 12 months (SCA6) |  |  |  | <0.001 |
| 5 months x 12 months (Control) |  |  |  | <0.001 |
| 7 months x 12 months (Control) |  |  |  | 0.003 |

**Table S5.** Statistics for impact of cerebellar Gq-DREADD stimulation on the performance of SCA6 84Q mice in the Morris water maze: repeated measures two-way ANOVA.

| Factor | df<sub>num</sub> | df<sub>den</sub> | F | p |
| --- | --- | --- | --- | --- |
| ***Escape Latency*** |  |  |  |  |
| Genotype | 2 | 22 | 3.821 | 0.038 |
| Time (SCA6 DREADD) | 9 | 54 | 20.887 | <0.001 |
| Time (SCA6 mCherry) | 9 | 72 | 16.160 | <0.001 |
| Time (Control mCherry) | 9 | 72 | 11.884 | <0.001 |
| Pairwise comparisons (p≤0.05 only): |  |  |  |  |
| SCA6 DREADD x SCA6 mCherry |  |  |  | 0.017 |
| Control mCherry x SCA6 mCherry |  |  |  | 0.049 |
| ***Distance moved*** |  |  |  |  |
| Genotype | 2 | 22 | 4.057 | 0.032 |
| Time (SCA6 DREADD) | 9 | 54 | 19.635 | <0.001 |
| Time (SCA6 mCherry) | 9 | 72 | 15.482 | <0.001 |
| Time (Control mCherry) | 9 | 72 | 10.187 | <0.001 |
| Pairwise comparisons (p≤0.05 only): |  |  |  |  |
| Control mCherry x SCA6 mCherry |  |  |  | 0.01 |
| ***Distance to Platform*** |  |  |  |  |
| Genotype | 2 | 22 | 4.921 | 0.017 |
| Time (SCA6 DREADD) | 9 | 54 | 23.635 | <0.001 |
| Time (SCA6 mCherry) | 9 | 72 | 12.613 | <0.001 |
| Time (Control mCherry) | 9 | 72 | 20.754 | <0.001 |
| Pairwise comparisons (p≤0.05 only): |  |  |  |  |
| SCA6 DREADD x SCA6 mCherry |  |  |  | 0.006 |
| Control mCherry x SCA6 mCherry |  |  |  | 0.048 |
| ***Velocity*** |  |  |  |  |
| Genotype | 2 | 22 | 0.921 | 0.413 |
| Time (SCA6 DREADD) | 9 | 54 | 1.361 | 0.229 |
| Time (SCA6 mCherry) | 9 | 72 | 2.933 | 0.005 |
| Time (Control mCherry) | 9 | 72 | 2.589 | 0.012 |

